# Proteomic and Lipidomic Plasma Evaluations Reveal Biomarkers for Domoic Acid Toxicosis in California Sea Lions

**DOI:** 10.1101/2024.05.06.592757

**Authors:** Amie M. Solosky, Iliana M. Claudio, Jessie R. Chappel, Kaylie I. Kirkwood-Donelson, Michael G. Janech, Alison M. Bland, Frances M.D. Gulland, Benjamin A. Neely, Erin S. Baker

**Affiliations:** Department of Chemistry, University of North Carolina at Chapel Hill, Chapel Hill, NC, USA; Department of Chemistry, North Carolina State University, Raleigh, NC, USA; Department of Biological Sciences, North Carolina State University, Raleigh, NC, USA; Immunity, Inflammation, and Disease Laboratory, National Institute of Environmental Health Sciences, Durham, NC, USA; Department of Biology, College of Charleston, Charleston, SC, USA; Wildlife Health Center, University of California, Davis, CA, USA; Chemical Sciences Division, National Institute of Standards and Technology, Charleston, SC, USA

**Keywords:** Domoic Acid, Proteomics, Lipidomics, California Sea Lion, Zalophus californianus, Ion Mobility Spectrometry, Multi-omics

## Abstract

Domoic acid is a neurotoxin secreted by the marine diatom genus, *Pseudo-nitzschia*, during toxic algal bloom events. California sea lions (*Zalophus californianus*) are exposed to domoic acid through ingestion of fish that feed on toxic diatoms, resulting in a domoic acid toxicosis (DAT), which can vary from mild to fatal. Sea lions with mild disease can be treated if toxicosis is detected early after exposure, therefore, rapid diagnosis of DAT is essential but also challenging. In this work, we performed multi-omics analyses, specifically proteomic and lipidomic, on blood samples from 31 California sea lions. Fourteen sea lions were diagnosed with DAT based on clinical signs and postmortem histological examination of brain tissue, and 17 had no evidence of DAT. Proteomic analyses revealed three apolipoproteins with statistically significant lower abundance in the DAT individuals compared to the non-DAT individuals. These proteins are known to transport lipids in the blood. Lipidomic analyses highlighted 29 lipid levels that were statistically different in the DAT versus non-DAT comparison, 28 of which were downregulated while only one was upregulated. Furthermore, of the 28 downregulated lipids, 15 were triglycerides, illustrating their connection with the perturbed apolipoproteins and showing their potential for use in rapid DAT diagnoses.

**SYNOPSIS:** Multi-omics evaluations reveal blood apolipoproteins and triglycerides are altered in domoic acid toxicosis in California sea lions.

**GRAPHIC ABSTRACT:** 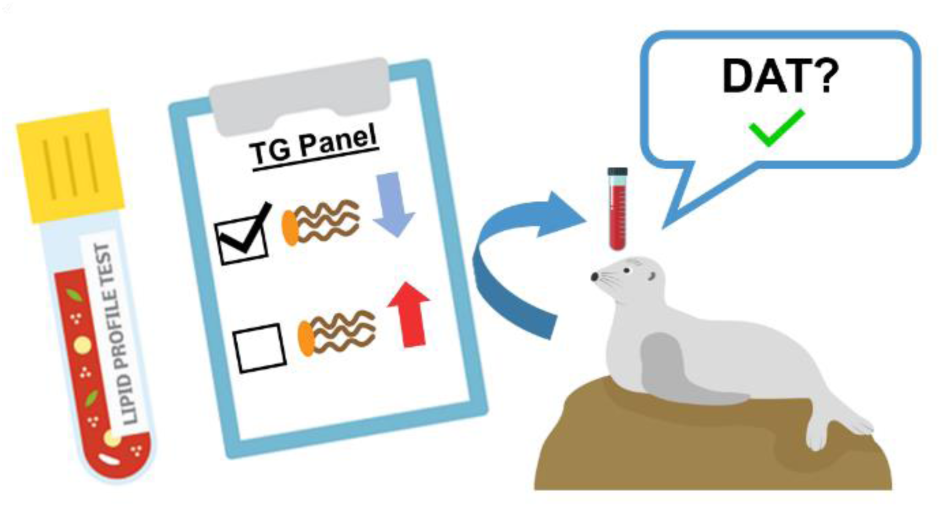

## INTRODUCTION

Domoic acid is a naturally occurring toxin produced by the pennate diatom, *Pseudo-nitzschia^1^*, a harmful algal bloom that is increasing in frequency and extent along the U.S. west coast.*^2^* In birds and mammals, domoic acid acts as an excitotoxin, affecting the brain and causing permanent short-term memory loss, seizures, brain damage, and death in certain situations.*^3, 4^* Specifically, exposure to domoic acid results in domoic acid toxicosis (DAT) in California sea lions (*Zalophus californianus)* and other animals such as seabirds and even humans.*^3,5,6^* More specifically, sea lions are exposed to domoic acid through anchovies and sardines in their diet that accumulate the toxin from eating and swimming in the algae (**Figure 1**).*^7^* Following exposure, domoic acid binds to glutamate receptors in the sea lion’s brain causing neurological dysfunction including seizures and ataxia.*^8^* Additionally, the toxin can bind to receptors in the heart, giving rise to cardiomyopathy and fatality.*^3^*

**Figure 1.**
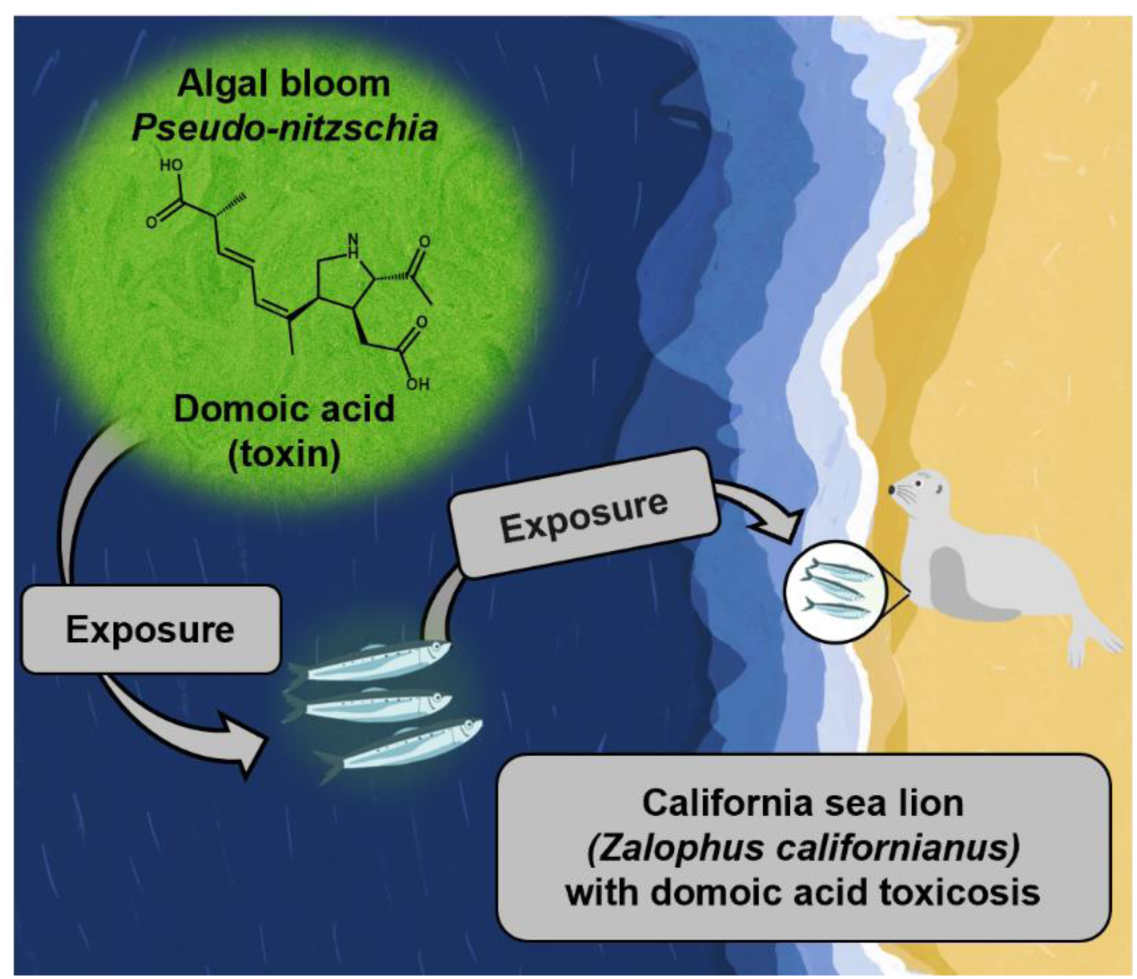
California sea lion domoic acid exposure routes. First, *Pseudo-nitzschia* secretes domoic acid during algal blooms. The domoic acid is then accumulated in fish due to their diet of zooplankton that feed on algal cells, or the diatoms themselves. Finally, sea lions are exposed when eating contaminated fish. Depending on the amount of domoic acid present, this exposure can cause toxicosis, which can be fatal.

DAT has been impacting sea lions for over 25 years, causing them to strand on beaches and need immediate medical attention.*^5,6^* In 1998, it was estimated that The Marine Mammal Center (Sausalito, CA) rescued 400 sea lions, with at least 70 experiencing seizures, head weaving, or even unresponsiveness (coma) from DAT.*^6,7^* Between 1998 and 2019, over 11,000 California sea lions have been admitted to TMMC and evaluated for DAT.*^6^* Additionally, the annual rate of DAT in sea lions is expected to continue rising based on the 2.1 % increase per year between 1998 and 2019. *^6^* More concerningly, the rapid diagnosis of DAT is challenging since domoic acid is rapidly metabolized and not detected in urine or blood greater than 24 hours post exposure.*^6^* Current diagnostic approaches include examining clinical signs such as behavioral changes (seizures, coma, ataxia), antemortem MRIs, blood eosinophil cell counts, and histological samples of the brain postmortem.*^6^* While these methods have been successful, they are not specific, and require highly trained technicians and anesthesiologists. Furthermore, some can only be performed postmortem.*^6^* Despite diagnosis being a challenge, treatment for DAT in sea lions does differ than other ailments, for example, an individual suffering from DAT may be administered anti-seizure medications as well as supportive subcutaneous fluids to induce flushing of the toxin out of the body.*^6^* Thus, evaluating potential molecular biomarkers that can be found in blood samples taken while the animal is alive and at early stages of DAT would facilitate effective treatment and reduce DAT-associated mortality.

Multi-omic analyses are a promising approach for DAT biomarker evaluation, as they reveal molecular changes, such as protein and lipid changes, specific to DAT over other ailments.*^9,10^* Proteomic analyses of California sea lion plasma and cerebral spinal fluid were performed in 2015, resulting in 20 potential protein biomarkers for DAT with seven proteins upregulated in sea lion cerebral spinal fluid and apolipoprotein E downregulated in plasma.*^11,12^* Here we performed another proteomic assessment of sea lion plasma for 31 California sea lions (14 DAT and 17 non-DAT) due to the recent release of a new sea lion genome and newer mass spectrometry instrumentation. Additionally, due to previous detection of alteration of apolipoprotein E,*^12^* we added lipidomic analyses to further evaluate molecular alterations in DAT and assess potential biomarkers for better diagnosis of California sea lions.

## MATERIALS AND METHODS

### Sea Lion Blood Sample Collection, Preparation, and Inclusion & Exclusion Criteria

The California sea lion plasma was collected between 2007 and 2012, near or at their time of admission to The Marine Mammal Center (TMMC) in Sausalito, CA, following rescue due to numerous illnesses*^12^*. The TMMC Internal Animal Care and Use Committee approved protocols for this study (number 2007–2), and the research was conducted under the National Fisheries Service MMPA permit numbers 932-1489-00 and 932-1905-00. Data analysis was approved under NIST Research Protections Office (RPO) approval MML-AR20-0001. After collection, blood was placed in Na-citrate tubes, centrifuged at 3,000 rpm for 10 min, and the plasma layer was transferred to a cryovial and stored at -80 °C within 2 h. Thirty-one California sea lion samples were selected for the proteomic and lipidomic analyses due to their DAT and non-DAT statuses as defined by Neely et al.*^12^* Of these, fourteen of the individuals were diagnosed with DAT and the seventeen remaining had other ailments such as leptospirosis or were suffering from malnutrition. The hematological and individual characteristics of each animal are provided in **Table S1** and **Table S2**.

### Chemicals and Reagents

All chemical reagents unless otherwise stated were purchased from ThermoFisher Scientific (Waltham, MA). For specific catalog numbers for all chemicals, reagents, and equipment, see **Table S3**.

### Proteomic Sample Preparation

Prior to the study, the 31 sea lion plasma samples were randomized and assigned an experimental key to avoid bias and minimize batch effects. Each sample was then thawed and centrifuged at 13 000 x *g*_n_ for 8 min at 4 °C. Based on prior work with California sea lion and other mammalian plasma, a protein concentration of approximately 50 μg/μL was assumed for estimating the amount of enzyme for digestion. Therefore, 2 μL (approximately 100 μg protein) of each sample was added to 78 μL of 50 mmol/L ammonium bicarbonate buffer in water in Lo-Bind microcentrifuge tubes (Eppendorf; Hamburg, Germany); these aliquots from each sample were briefly mixed by vortexing and immediately used for digestion. Simultaneously, a pool was generated using 5 μL aliquots from each sample. The pool was digested in parallel to the individual samples and later used as a technical replicate during the LC-MS/MS analysis (**Figure 2A**). Proteins were reduced with 9 μL of 90 mmol/L DL-Dithiothreitol (DTT; final concentration of 9 mM) at 60 °C for 30 min, then cooled and alkylated with 9 μL of 200 mmol/L 2-chloroacetamide (CAA; final concentration of 18 mM) at room temperature in the dark for 30 min. An additional 100 μL of 50 mmol/L ammonium bicarbonate was added prior to enzymatic digestion. First, LysC (Thermo Scientific; Waltham, MA) was added at a 1:100 mass ratio (LysC:total protein) ratio, using 2 μL of a 0.5 μg/μL solution, and incubated at 37 °C for 1 h. Next, trypsin (Pierce Trypsin Protease, Thermo Scientific; Waltham, MA) was added at a 1:20 mass ratio (trypsin:total protein) using 5 μL of 1 μg/μL solution. Samples were incubated at 37 °C for 4 h, after which the reaction was halted by adding 5 μL of 4 % formic acid (volume fraction). Peptide samples were purified for LC-MS/MS using Pierce C18 spin columns (8 mg C18 resin) according to the manufacturer’s instructions. Briefly, 0.5 % trifluoroacetic acid (volume fraction) in 5 % acetonitrile (volume fraction) was used for wash steps, while 60 μL of 70 % acetonitrile (volume fraction) was used for elution. The samples were reduced to dryness in a vacuum centrifuge at low heat and stored at -20°C. Samples were then reconstituted with 100 μL of 0.1 % formic acid (volume fraction), vortexed for 10 min, and the Pierce quantitative fluorometric peptide assay was used to determine sample peptide concentrations with a BioTek Synergy HT plate reader (Agilent Technologies; Santa Clara, CA).

**Figure 2.**
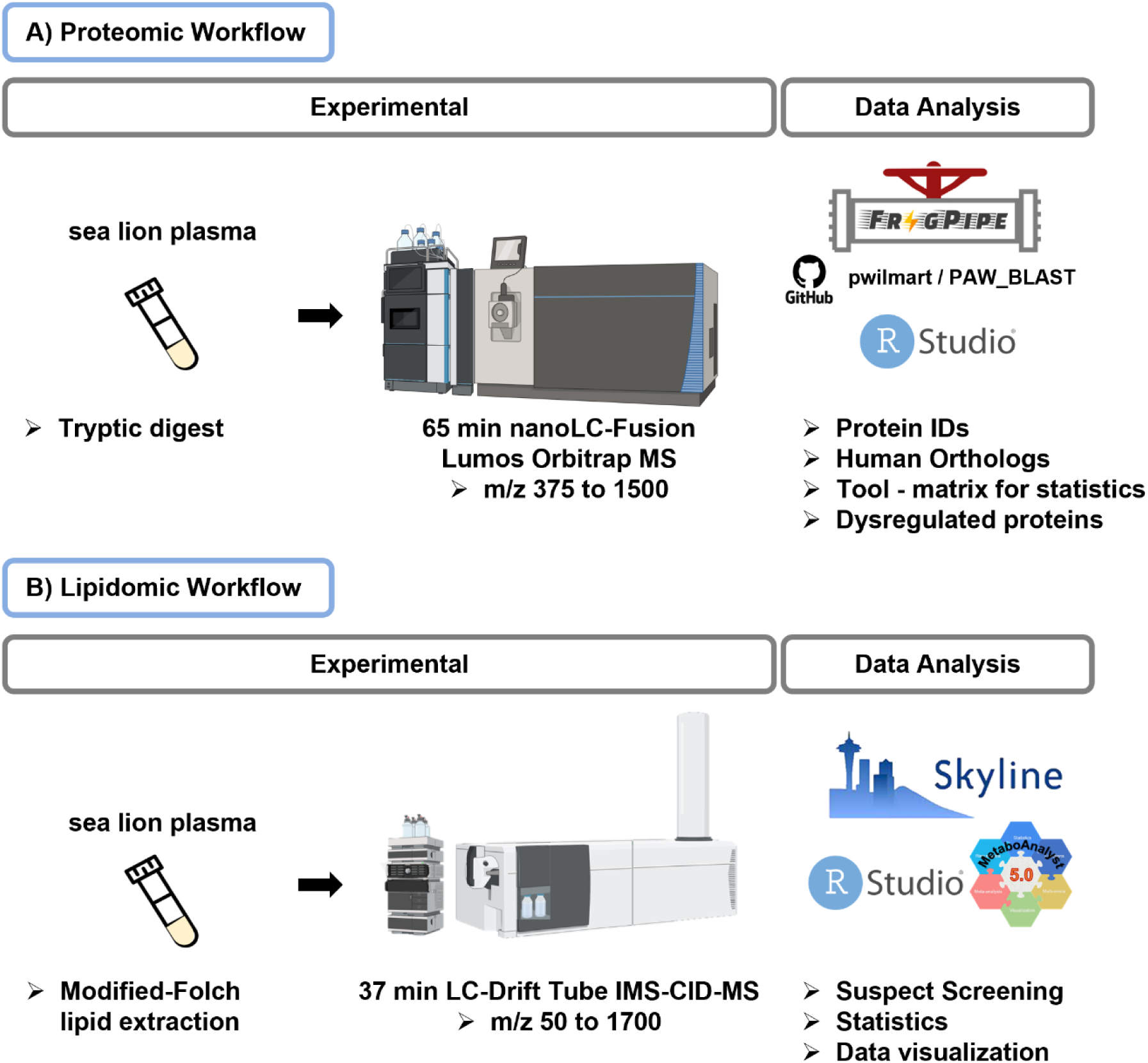
Proteomic and lipidomic sample workflows. **A)** For the proteomic workflow, a tryptic digest of sea lion was separated with nanoflow liquid chromatography and data acquired with a Fusion Lumos orbitrap mass spectrometer. Data was searched with FragPipe and analyzed in R. **B)** The lipidomic workflow consisted of a modified-Folch lipid extraction of the plasma, followed by LC-IMS-CID-MS analysis. Lipidomic data sets were processed using Skyline software with in-house libraries containing over 1,000 lipids with LC retention time, CCS, and *m/z* values for each. Statistics and data visualization were performed using MetaboAnalyst 6.0 and RStudio. The figure was created with BioRender.

### Proteomic nanoLC-MS/MS Instrumentation

Peptide mixtures in 0.1 % formic acid (volume fraction) were analyzed using an UltiMate 3000 nanoLC coupled to a Fusion Lumos Orbitrap mass spectrometer (Thermo Fisher Scientific; Waltham, MA). Peptide mixtures were loaded onto a PepMap 100 C18 trap column (75 µm id x 2 cm length; Thermo Fisher Scientific; Waltham, MA) at 3 µL/min for 10 min with 2 % acetonitrile (volume fraction) and 0.05 % trifluoroacetic acid (volume fraction) followed by separation on an Acclaim PepMap RSLC 2 µm C18 column (75µm id x 25 cm length; Thermo Fisher Scientific; Waltham, MA) at 40 °C. A randomized sample order and injection volumes were determined for 1 μg loading (between 2.8 μL and 7.7 μL). Peptides were separated along a 65 min gradient with mobile phase A (MPA) consisting of 0.1 % formic acid (volume fraction) in water and mobile phase B (MPB) being 80 % acetonitrile (volume fraction) and 0.08 % formic acid (volume fraction) in water. A two-step gradient was used, beginning with 5 % to 30 % MPB over 50 min followed by 10 min ramp to 45 % MPB, and lastly to 95 % MPB over 5 min, and held at 95 % MPB for 5 min, all at a flow rate of 300 nL/min. The Fusion Lumos was operated in positive polarity with 30 % RF lens, data-dependent mode (topN, 3 sec cycle time) with a dynamic exclusion of 60 s (with 10 ppm error). Full scan resolution using the orbitrap was set at 60 000, the mass range was *m/z* 375 to 1500. The full scan ion target value was 4.0 x 10*^5^* allowing a maximum injection time of 50 ms. Monoisotopic peak determination was used, specifying peptides and an intensity threshold of 2.5 x 10^4^ was used for precursor selection, including charge states 2 to 6. Data-dependent fragmentation was performed using higher-energy collisional dissociation (HCD) at a normalized collision energy of 32 with quadrupole isolation at *m/z* 1.3 width. The fragment scan resolution using the orbitrap was set at 15 000 for *m/z* 100 as the first mass, ion target value of 2.0 x 10^5^, and 30 ms maximum injection time. The MS1 data was collected as profile data, while the MS2 data was collected as centroid data. The function, inject all ions for parallelizable time, was not used in the analysis. The raw data and tissue specific search results along with all databases used are available at the ProteomeXchange Consortium*^13^* via the PRIDE partner repository with the dataset identifiers PXD020366 and 10.6019/PXD020366.*^14^*

### Protein Identification and Label-free Quantification

The data-dependent acquisition data was searched with FragPipe (v20.0), a software package that includes the MSFragger search algorithm (v3.8),*^15^* Philosopher protein inference model (v5.0.0),*^16^* and an IonQuant label-free quantification algorithm (v1.9.8).*^17^* The parameter file used (which can be viewed and loaded into FragPipe) is also available as supplemental **File S1**. The search parameters were chosen for trypsin and LysC, and it is known that non-specific enzyme activity occurs in biofluids. Broadly, the parameters included semitryptic cleavage with a strict trypsin cleavage rule (cut after KR) and allowing for two mis-cleavages. The precursor and fragment mass tolerances were initially set to 20 ppm for precursor and fragment mass tolerance, though MSFragger runs a mass calibration and optimization step to set these tolerances automatically. Oxidized methionine, protein n-term acetylation, and peptide n-term pyroGlu were allowed as variable modifications and carbamidomethylation of cysteine was a fixed modification. The LFQ-MBR (label-free quantification match between runs) workflow was used, which is IonQuant-based MS1 quantification with match between runs. The built-in FragPipe function of fasta selection was used to select the UniProtKB *Zalophus californianus* (California sea lion; taxon ID 9704) proteome UP000515165 (47 492 sequences; retrieved 8 August 2023). This proteome is based on an updated NCBI RefSeq annotation that used the 2020 Vertebrate Genome Project Genome mZalCal1.pri.v2 assembly, but still used the underlying RNAseq data previously deposited*^18^*. Using FragPipe, 118 contaminant sequences (such as human keratin or porcine trypsin) were added, and finally the fasta was concatenated with reverse entries (95,230 entries total).

### Protein Ortholog Mapping

The UniProtKB *Zalophus californianus* (California sealion; taxon ID 9704) proteome UP000515165 contained mostly gene symbol IDs, therefore there was not a drastic need to perform orthology mapping on all 485 initially identified proteins (note some without quantification, see below). Instead, only 71 of the identified proteins were missing gene symbols and had locus numbers (ex. LOC118356106 for serum amyloid protein A). Human orthologs were assigned to these 71 proteins using a series of python scripts (Anaconda v2023.07; conda v23.5.2; Python v3.8.17) from Github on the following two repositories: pwilmart/PAW_BLAST and pwilmart/annotations, retrieved October 12, 2021. First, the 71 identifiers in question were used to create a subset fasta (make_subset_DB_from_list_3.py), which was then searched against a human fasta [UniProtKB *Homo sapiens* (Human; taxon ID 9606) canonical proteome UP000005640; 2023_03 release] using the db_to_db_blaster.py script, which used a local installation of BLAST+ 2.11.0. This list was manually confirmed by comparing sea lion to human ortholog protein names, and when needed, running BLAST with sea lion entries against all mammals. Incorrect assignments can happen when a protein is not present in humans (e.g., beta-lactoglobulin 1; UniProtKB A0A6P9EXQ1), or if the human BLAST hit disagrees with the original annotation, in which case the original annotation is preserved (e.g., sea lion haptoglobin was matched to human haptoglobin-related protein, but we kept the original name of haptoglobin and the HP symbol). Once all 485 identified sea lion proteins had gene symbols, any proteins with all zeros by MaxLFQ intensity were removed (this can occur when a tandem mass spectrum is identified in FragPipe, but a quantity not calculated). Next, any remaining duplicate gene symbol entries were summed together into a single nonredundant entry. This resulted in 245 proteins identified experiment-wide, which were used for statistical testing (**Table S5**).

### Proteomic Statistical Data Analysis

A two-sided Wilcoxon rank sum test (equivalent to the Mann-Whitney U-test) was used to detect statistically significant proteins. This was performed with R (v4.2.0) and the dplyr package (v1.1.0) using the *wilcox.test* function and p-value correction with the Benjamini and Hochberg method, using the *p.adjust* function specifying ‘BH’.*^19^* Proteins with adjusted p < 0.05 (*i.e.,* FDR < 0.05) were deemed significantly different.*^19^* Since zeros cannot be used to calculate fold-change, all zeros were replaced with ½ the lowest observed non-zero value for each given protein. This imputation did not affect rank sum statistics but is used solely to provide a reference for protein change between groups.

### Lipidomic Sample Preparation

For the lipidomic analyses, lipids from the 31 individual plasma samples, as well as four pooled samples, were extracted using a modified-Folch extraction.*^20^* This lipid extraction method was previously described and validated by Kirkwood et al., using NIST SRM 1950 (metabolites in frozen human plasma) as a quality control to compare with other extraction methods.*^22^* The modified-Folch extraction resulted in a comparable amount of lipids, as well as category compositions, as various other extraction methods performed by the LIPID MAPS consortium.*^22^* See **Figure 2B** for the lipidomic workflow used in the present study. Briefly, 50 μL of plasma was transferred into a 1.7 mL Sorensen SafeSeal™ Microcentrifuge tube (Waltham, MA) with 600 μL of 2:1 (volume ratio) chloroform/LC-MS grade methanol at -20 °C. Each sample was vortexed for 30 s at 1200 rpm using a Digital Vortex Mixer (Thermo Scientific; Waltham, MA) and then 150 μL of LC-MS grade water was added, prior to mixing again for 30 s. The samples were then left at room temperature for 5 min to allow adequate time for phase separation, centrifuged at 12 000 rpm for 10 min at 4 °C for further phase separation, and immediately placed on ice to prevent degradation.*^21^* Since this liquid/liquid extraction results in both a top (aqueous) and bottom (organic) extraction layer, 100 μL of the bottom layer was aliquoted into a new tube and dried down using a SpeedVac vacuum concentrator (ThermoFisher Scientific; Waltham, MA). Following dry down, the lipid extract was immediately reconstituted in 10 μL of chloroform and 190 μL of LC-MS grade methanol. The samples were stored in 2-mL amber glass LC autosampler vials (Agilent Technologies; Santa Clara, CA) at -20 °C until analysis (less than 1 week). The lipidomic pooled quality control sample was made by combining equal amounts of each individual sample into a new 1.7 mL Sorensen SafeSeal™ Microcentrifuge tube (Sorensen Bioscience, Inc.; Murray, UT). The pool was then vortexed for 30 s and underwent the same modified-Folch extraction steps as above in parallel with the rest of the sample set. The final QC sample was split into four vials to provide technical replicates for the LC-IMS-CID-MS analyses.

### Lipidomic LC-IMS-CID-MS Instrumentation

Lipidomic analyses of the 31 plasma samples were performed using a previously described untargeted platform combining liquid chromatography, ion mobility spectrometry, collision-induced dissociation, and mass spectrometry (LC-IMS-CID-MS).*^22^* **Figure 2B** illustrates the lipidomic workflow. These separations were performed on an Agilent 1290 Infinity UPLC system (Agilent Technologies; Santa Clara, CA) coupled to an Agilent 6560 IM-QTOF platform (Agilent Technologies; Santa Clara, CA) with a commercial gas kit and a MKS Instruments precision flow controller (Andover, MA).*^22^* Prior to all lipidomic studies, Agilent ESI-L Low Concentration tuning mix solution (Agilent Technologies; Santa Clara, CA) was injected in the 6560 IM-QTOF platform to verify mass accuracy and collision cross section (CCS) values of the platform. Brain total lipid extract standard (Avanti Polar Lipids; Alabaster, AL) was run prior to samples to assess lipidomic separations and quality. Then, 10 µL of each individual plasma lipid extract was injected onto a Waters Acquity UPLC CSH^TM^ C18 column (Milford, MA) (3.0 mm x 150 mm x 1.7 μm particle size), where MPA was comprised of 40:60 (volume ratio) LC-MS grade ACN/H_2_O and 10 mmol/L ammonium acetate and MPB consisted of 90:10 (volume ratio) LC-MS grade IPA/ACN with 10 mmol/L ammonium acetate. The 34 min LC gradient had a constant flow rate of 0.25 mL/min and time to MPB ratio as follows [time (min):MPB (%; volume fraction to MPA)): 0:40, 2:50, 3:60, 12:70, 15:75, 17:78, 19:85, 22:92, 25:99, and 34:99]. The gradient was followed by a 3 min column wash step as follows [flow rate (mL/min): time (min):MPB (%; volume fraction to MPA)): 0.30:34.5:40, 0.30:35:99, 0.30:35.5:99, 0.35:36:40, 0.30:37:40, and 0.25:38:40]. For the IMS-MS analyses, the Agilent Jet Stream ESI source was utilized, and the platform was operated in both negative and positive ionization modes. For the IMS analyses, the funnel trap fill time was set to 10 ms with a release time of 400 μs. Following drift time separation, the CID analysis consisted of a data-independent approach where lipid ions were fragmented using a CID ramp based on the precursor ion’s drift time (**Table S4)**. Finally, a quadrupole time-of-flight (QTOF) mass spectrometer evaluated ion from 50 to 1700 *m/z*, and a .d file was collected containing the multidimensional LC, IMS, CID, and MS information.

The .d LC-IMS-CID-MS data files were then single-field calibrated to calculate CCS values for all lipids in the samples using the Agilent MassHunter Workstation Software IM-MS Browser Version B.10.00 and the previously mentioned Agilent tuning mix.*^23^* All raw data files were evaluated using Skyline software, an open source, non-vendor specific software to process various types of mass spectrometry data (MacCoss Lab Software, v. 22.2).*^24,25^* In Skyline, lipids were annotated based on how closely the extracted ion chromatograms matched our in-house spectral lipid library, which contained 877 lipids and is publicly available at https://tarheels.live/bakerlab/databases..*^22,26^* The parameters manually assessed were retention time index, *m/z* values with a mass error of less than 5 ppm, and CCS values within the resolving power window of 50.*^26^* Peaks were additionally validated by precursor isotope distribution and fragmentation spectra. The raw data is available on Panorama at https://panoramaweb.org/sealionlipids.url.

### Lipidomic Statistical Analysis

The list of lipid identifications and their corresponding peak areas in the samples from Skyline were then normalized via total ion current and log_2_ transformed. The normalized and transformed peak areas were imported into MetaboAnalyst 6.0 to assist in data visualization using hierarchical clustering (based on Euclidean distances) in the form of a heatmap to assess differences between the DAT and non-DAT sea lions.*^27^* Additional statistical analyses were also performed using RStudio (R version 4.3.1) including the following packages: dplyr and Lipid Mini-On (rodin).*^19,28^* In the same fashion as the proteomic statistical analysis, the lipidomic statistical comparisons included a two-sided Wilcoxon rank sum test and fold change comparisons. Additionally, volcano plots and bar charts were used for data visualization.

## RESULTS & DISCUSSION

### Proteomic Analyses

In the proteomic analyses of the 31 plasma samples, 245 proteins were identified from 9079 peptides (**Table S5**) and statistical assessments of the DAT versus non-DAT groups found 31 proteins were statistically significant (Benjamini-Hochberg adjusted p-value < 0.05, **Table S7**). Of these proteins, three were apolipoproteins (APO): APOC3, APOE, and APOB (noted with * in **Figure 3**) and an additional eight apolipoproteins were also detected with specific trends. Since APOE was observed to be of statistical significance and decreased in this sample previously using 2D-gel, its results were not surprising.*^12^* However, APOC3 downregulation in DAT individuals was of great interest as its high levels have been linked to hypertriglyceridemia in humans, indicating a potential for hypotriglyceridemia in the DAT group.*^29^* Downregulation of APOB has also been linked with decreased triglyceride availability in humans, and as the protein plays a role in lipid metabolism, this further illustrates the need for a lipidomics assessment.*^30^* Additionally, of the 11 apolipoproteins detected in this study while eight were not statistically significant they did have interesting trends. For example, APOC1 and APOC4 had lower abundances in the DAT group and are thought to be involved in triglyceride metabolism, although the function of endogenous APOC4 is not completely known.*^31,32^* APOC2 was also not statistically significant but trended higher in the DAT group, and high APOC2 levels have been associated with downregulation of plasma TGs in humans.*^32^* Since this study size is small, the trends in these non-significant apolipoproteins are also notable.

**Figure 3.**
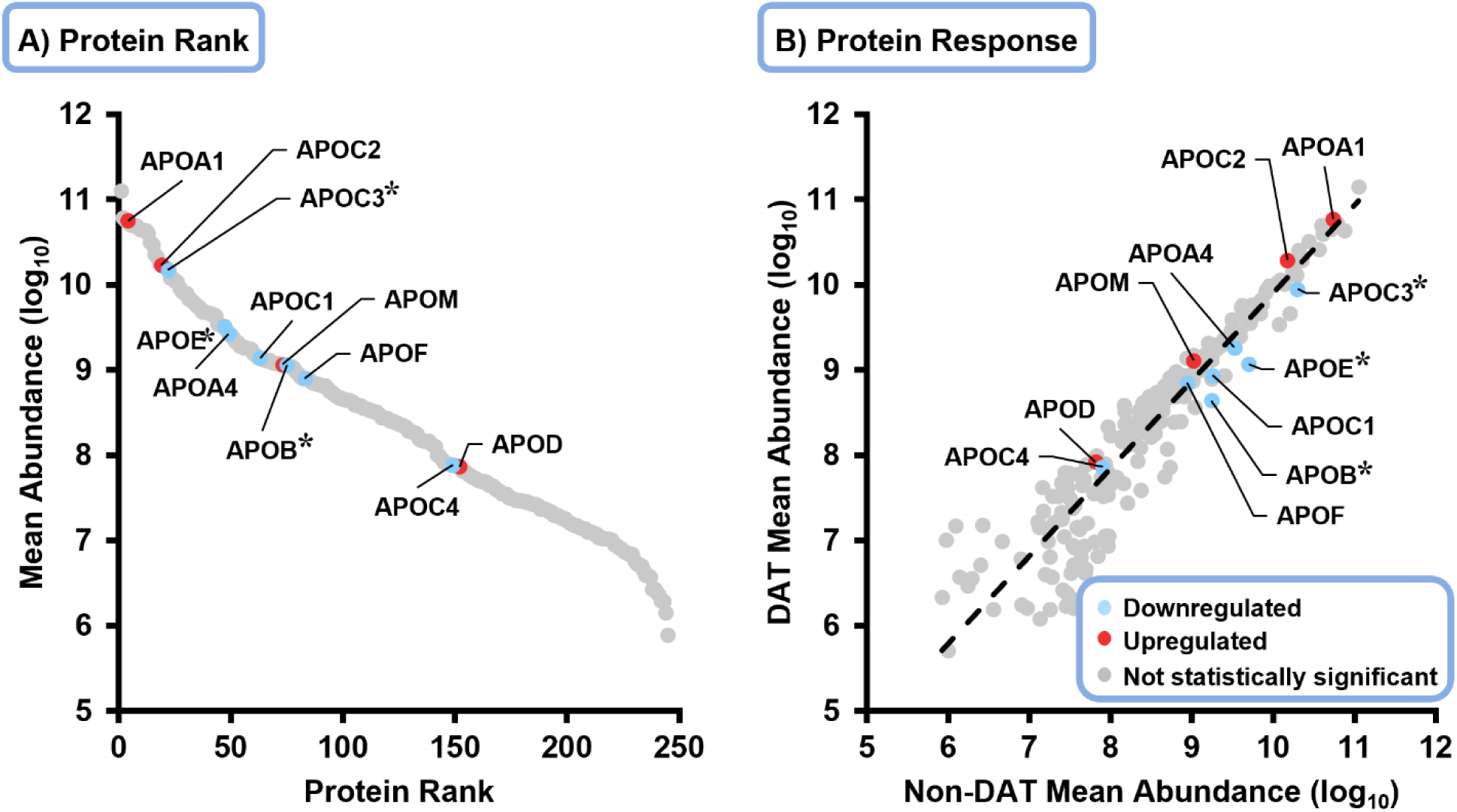
Eleven apolipoproteins detected in the sea lion proteomic analyses and illustrated in **A)** rank abundance and **B)** protein response graphs. Apolipoproteins are highlighted in either red or blue, indicating up (red) or downregulation (blue), while all other detected proteins are shown in grey. The dashed line in **B)** represents a 1:1 reference, where proteins above are upregulated and below are downregulated. The statistically significant apolipoproteins are noted with asterisks.

Of the other 28 proteins found to be statistically significant altered in DAT in the study, other interesting trends were observed and angiopoietin-like 3 (ANGPTL3), hepatocyte growth factor activator (HGFA), coagulation factor XII, and CD5 antigen-like protein (CD5L) were of particular interest. For example, ANGPTL3 had the lowest log_2_ fold change (approximately -4) out of all the significant proteins and is involved in blood TG regulation.*^33^* HGFA and coagulation factor XII, which were higher in DAT individuals, are both proteases thought to activate hepatocyte growth factor signaling,*^34^* a signaling pathway activated when there is injury to tissues.*^34^* Furthermore, coagulation factor XII has been linked to pro-inflammatory cytokines in antigen-presenting cells in atherosclerosis-prone mice, indicating an increased immune response,*^35^* which could help explain why sea lions with DAT may suffer from cardiomyopathy. Lastly, CD5L, which was downregulated in the DAT individuals, is involved in lipid homeostasis and immunomodulation. Additionally, it has been linked to mediate autoimmunity by altering fatty acid composition and inhibiting cholesterol biosynthesis.*^36,37^* Thus, in addition to the apolipoproteins, all four of these proteins of interest have been linked to lipids and further support our hypothesis that lipids are dysregulated in individuals with DAT.

### Lipidomics Analyses

In the lipidomic analyses of the 31 sea lion plasma samples, 331 unique lipids in five lipid categories (fatty acyl, sterol, glycerolipid, glycerophospholipid, and sphingolipid) were identified at a Schymanski confidence level of 2 or 3.*^22,26,38^* Of the identified lipids, 29 were found to be statistically significant in the DAT versus non-DAT comparison, with 28 downregulated and 1 upregulated (**Table S8**). Moreover, while the statistically significant lipids span four out of the five lipid categories assessed, 15 of the 29 were triglycerides (TGs), illustrating their potential link to DAT and its apolipoprotein dysregulation. In general, the glycerolipids (TGs and diglycerides (DGs)) were remarkably enriched, making up just 15 % of all 331 detected lipids and 55 % of the significant. Furthermore, all but one of the significant lipids were lower in the DAT sea lion plasma, including all TGs as shown in the volcano plot in **Figure 4**. The single upregulated significant lipid was a phosphatidylinositol species (PI(18:0_20:3)). Interestingly, phosphatidylinositols are precursors for DGs, which is another glycerolipid like TGs but with one less fatty acyl chain.*^39^* This is worth noting because one diglyceride, DG(18:1_18:1), was found to be downregulated and statistically significant in the DAT individuals, which indicates that a lack of DG(18:1_18:1) synthesis may a consequence when a higher abundance of PI(18:0_20:3) is observed due to the PI not being converted to a DG, but requires more investigation.

**Figure 4.**
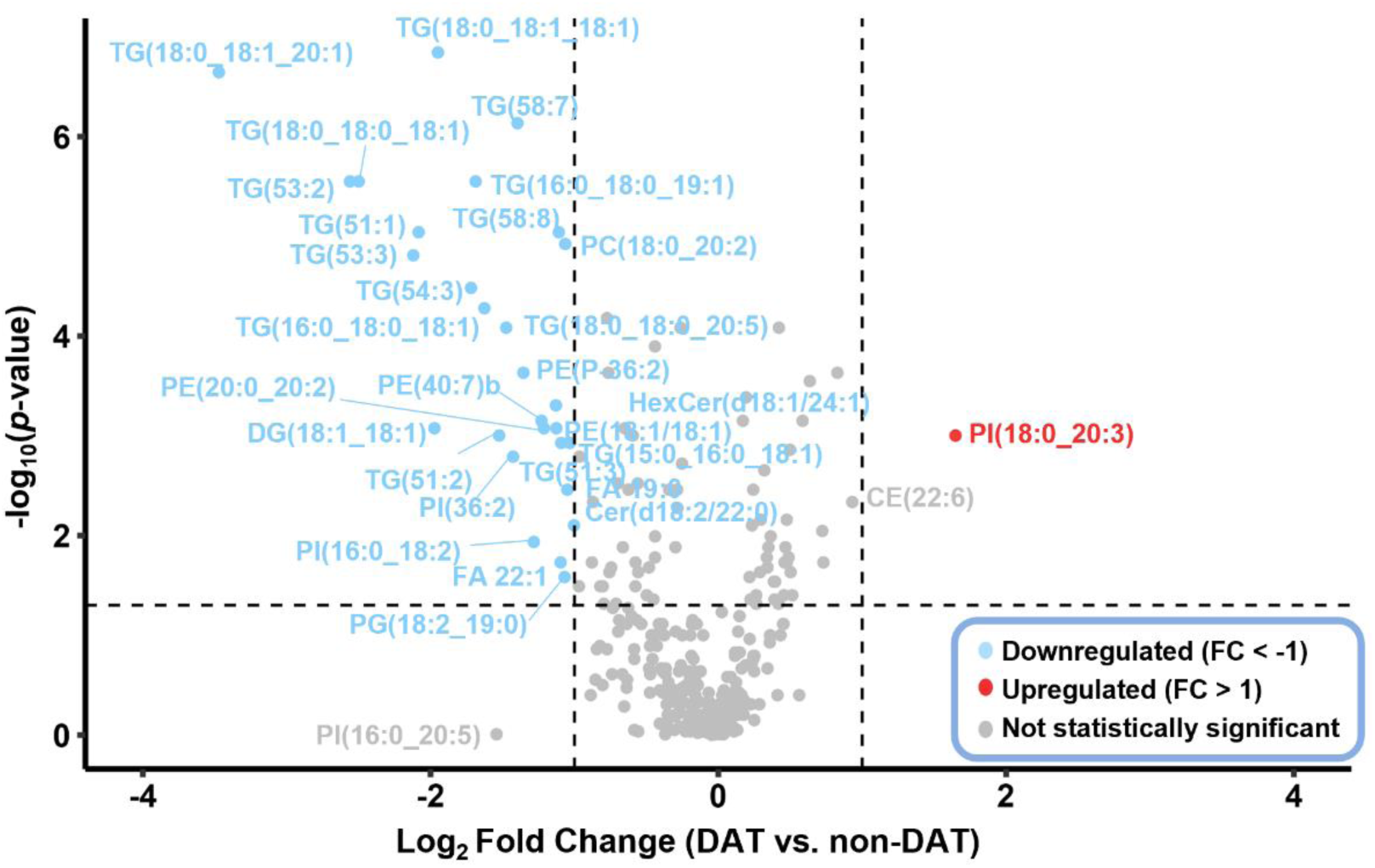
Lipids detected and dysregulated in the DAT versus non-DAT California sea lion plasma comparison. Blue data points show downregulated lipids, red show upregulated, and grey show those not statistically significant in the comparison. Of the significant lipids, 50 % are triglycerides. The full list of significant lipids is included in **Table S8**.

To further evaluate the lipidomic changes in DAT individuals, the significant lipids were assessed hierarchical clustering in Metaboanalyst 6.0*^27^*(**Figure 5**). Here, hierarchical clustering illustrated that the DAT and non-DAT individuals clustered, and similar trends were observed in all glycerolipids, including the 15 TGs and 1 DG, plus 1 glycerophospholipid, PE(18:1/18:1). Specifically, these lipids exhibit downregulation in the DAT samples, suggesting a link between DAT and decreased TGs. Furthermore, many of the trending downregulated proteins are involved with lipoprotein lipase, such as APOA1 and APOC2, which is responsible for the regulation of TG degradation.*^32^* This further couples with the observed proteomic and lipidomic changes observed in DAT. Interestingly, this was an unsupervised heatmap, thus the sample groups clustered together without supervision. Consequentially, this clustering revealed another trend: four non-DAT individuals had similar molecular compositions as those in the DAT group. Two of those individuals were diagnosed with leptospirosis, another condition that is common among sea lions.*^6^* One of those had trauma to the eye as well, which is worth noting because of the two that did not have leptospirosis in that clustered group, one of them had flipper trauma. The last individual was malnourished, which might explain why their abundance of TGs was lower than the rest of the non-DAT individuals, since they may have had to rely on lipid energy stores to survive. The malnourished individual is the left most sample in the heatmap, indicating that it clustered more with the non-DAT samples. This illustrates a potential continuum between TG abundance and DAT but requires further research to state conclusively.

**Figure 5.**
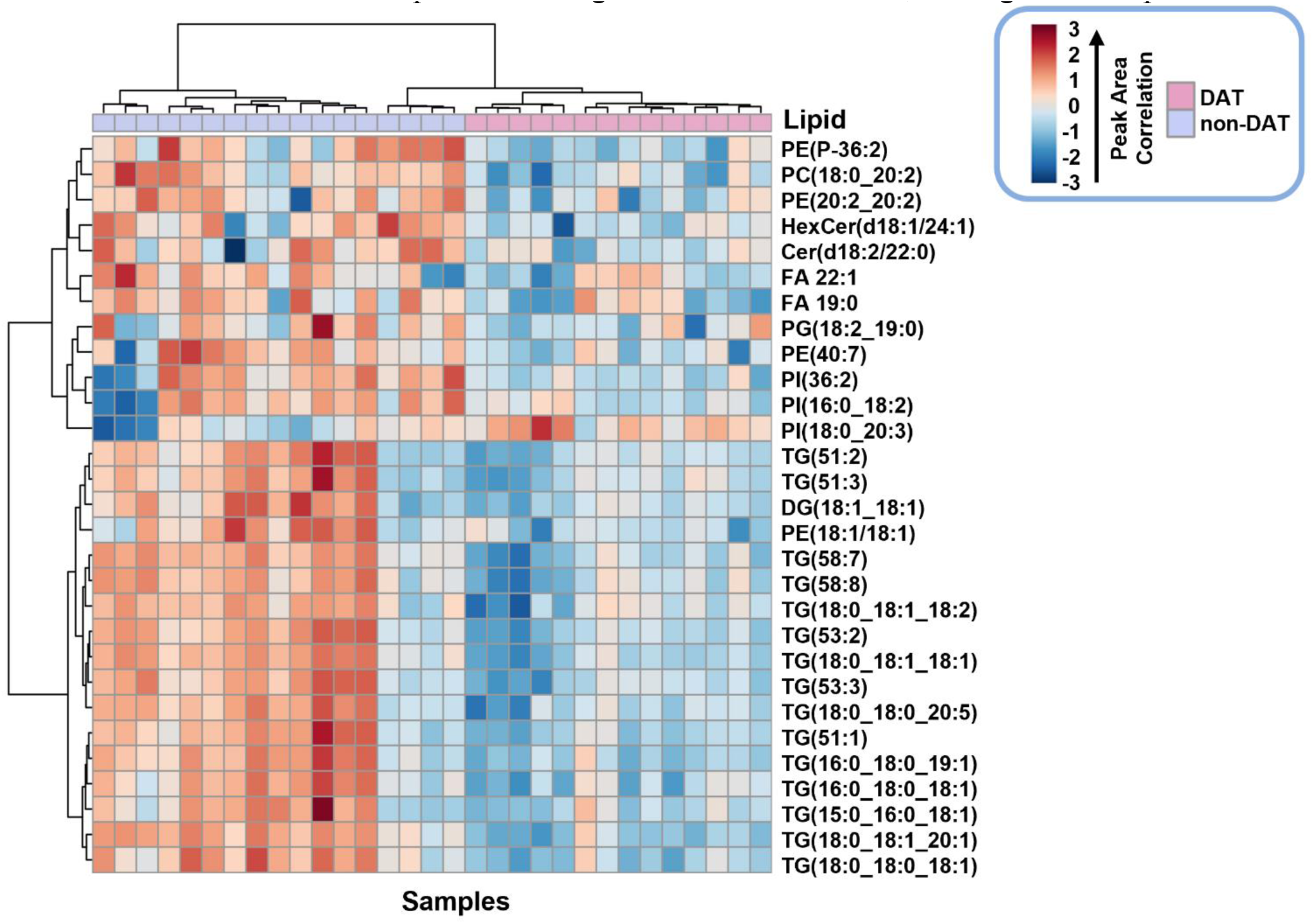
Hierarchical clustering of the California sea lion plasma samples for significant lipids detected in both positive and negative mode. Only the significant lipids are shown in each mode. Lipids with higher abundances are shown in red and those with less abundant in blue. The Euclidean distances were calculated and clustered using the Ward method.

To further probe lipid structure, trends in the degrees of unsaturation and fatty acyl chain lengths were investigated to find patterns in the types of lipids found to be potential biomarkers (**Figure 6**). Specifically, 58 % of the statistically significant lipids were found to have at least one degree of unsaturation and 48 % had at least one fatty acyl chain with 18 carbons (considered long chain fatty acids (LCFA)). Furthermore, diunsaturated lipids represented 9 % of the total lipids detected but increased in representation to almost 18 % of the lipids found to be statistically significant in DAT individuals. This 18-carbon fatty acyl chain and diunsaturation trend is further supported by the volcano plot in **Figure 4**. Specifically, TG(18:0_18:1_20:1) which has two 18 carbon chains (all three LCFAs) and 2 double bonds has the lowest *p*-value (of the lipids with fatty acyl chain assignments) and highest magnitude log_2_ fold change. Other statistically significant TGs were also observed with 18 carbon chains and unsaturation (**Figure 5**), illustrating how TGs could be quite informative of DAT in sea lions. The 15 statistically significant TGs downregulated in the DAT group also support the idea that a simple TG blood panel assay could be developed for a more accessible DAT evaluation platform, such as ultraviolet-visible spectroscopy. This would enable rapid and affordable diagnostic testing for DAT and be extremely beneficial for the veterinarians and sea lions.

**Figure 6.**
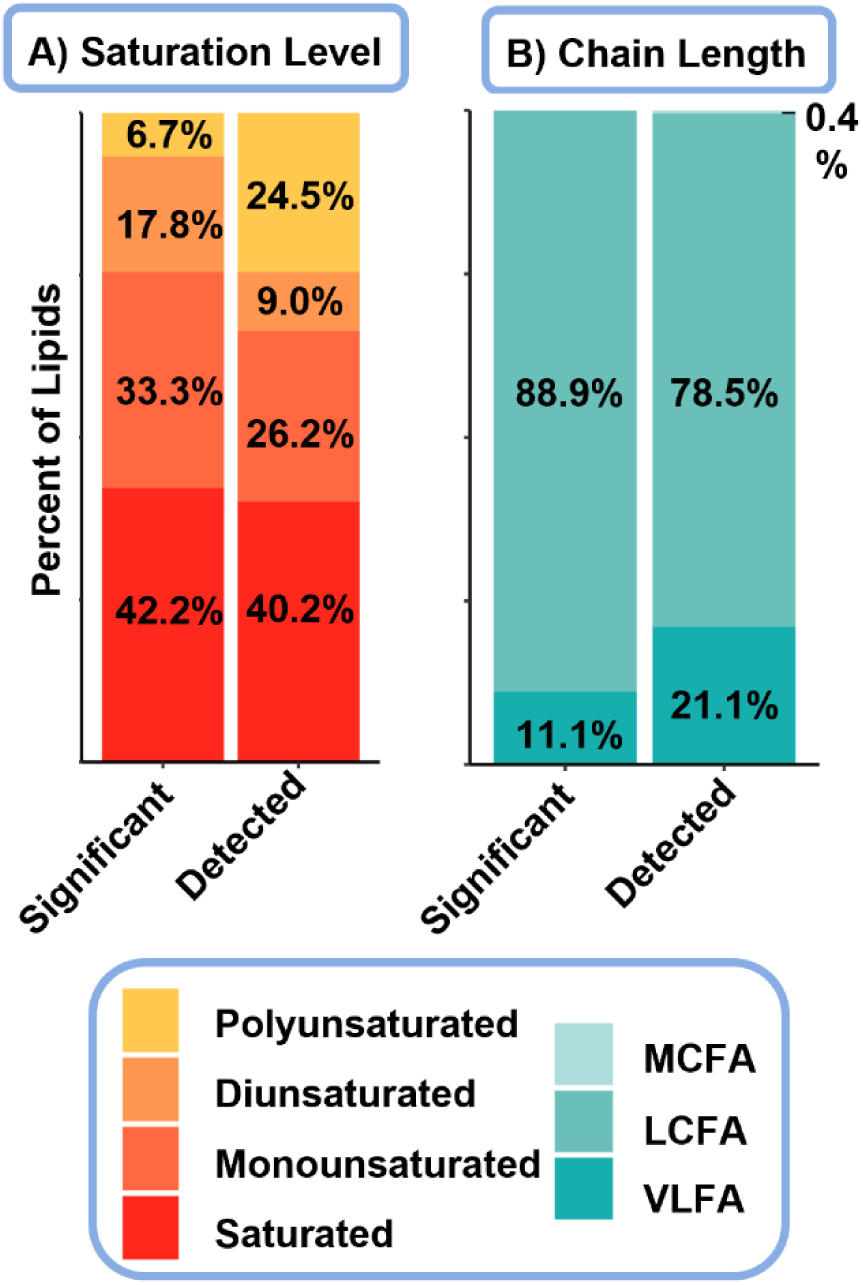
Saturation and chain length trends for detected and significant lipids in the DAT California sea lion plasma. **A)** The number of double bonds present in the fatty acyl chains of the significant and detected lipids, and **B)** variations in long chain fatty acids (13-20 carbons) or very long chain fatty acids (greater than 20 carbons). MCFA = medium chain fatty acids (6-12 carbons), LCFA = long chain fatty acids, and VLFA = very long chain fatty acids.

Although apolipoprotein and TG candidate biomarkers were determined in this work, this study does not go without limitations. First, the sample size of the study was only n = 31. This is a small group compared to the population of sea lions, especially considering the DAT group consisted of less than half of the total, n = 14. In addition to the study size, no healthy control group was present, as the non-DAT group had other ailments such as leptospirosis and starvation.*^12^*

However, healthy controls would be less useful for this study, as molecular markers for DAT are specifically needed for diagnosis and the treatment of sick animals presenting with a specific set of symptoms. Thus, future work using a larger sample size with more illnesses represented in the non-DAT group as well as true healthy controls would be important for confirming the results found in this study.

## CONCLUSIONS

In this study, multi-omic analyses were conducted on California sea lion plasma to identify potential biomarkers as diagnostic tools for DAT. The proteomic analyses confirmed APOE as a DAT plasma biomarker from a previous study*^12^* and showcased 2 other apolipoproteins as statistically significant and downregulated in the DAT sea lion plasma. In addition, 8 other apolipoproteins were noted as having specific trends in the analyses but were not noted as statistically significant, potentially because of the small sample size.*^12^* Lipidomic analyses of the plasma further illustrated the effects of apolipoprotein dysregulation in 15 TGs, which are associated with these proteins in the literature and were found to be less abundant in the sea lions with DAT as well as four non-DAT with similar symptoms. Since the TGs made up over 50 % of the significant lipids, they also provide a molecular class of interest that could be assayed (either with LC-MS or a less complex, but robust instrument such as UV-Vis) by veterinarians in conjunction with clinical signs such as seizures to provide a rapid diagnostic assessment. Although a larger study would help validate our findings, the proteins and lipids noted to differentiate DAT from non-DAT animals are extremely important for both understanding DAT molecular perturbations and creating potential diagnostic and treatment strategies.

## ACKNOWLEDGEMENTS

This work was funded by grants from the National Institute of Environmental Health Sciences (P42 ES027704), the National Institute of General Medical Sciences (R01 GM141277 and RM1 GM145416), and a cooperative agreement with the Environmental Protection Agency (STAR RD 84003201). This research was supported in part by the Intramural Research Program of the NIH (ZIC ES103363). The views expressed in this manuscript do not reflect those of the funding agencies. Identification of certain commercial equipment, instruments, software, or materials does not imply recommendation or endorsement by the National Institute of Standards and Technology, nor does it imply that the products identified are necessarily the best available for the purpose. We wish to thank Jackson Eberwein for his technical assistance related to sample preparation for proteomic analysis.

